# Parkinson’s disease and bacteriophages as its overlooked contributors

**DOI:** 10.1101/305896

**Authors:** George Tetz, Stuart M. Brown, Yuhan Hao, Victor Tetz

**Author notes:** Correspondence and requests for materials should be addressed to G.V.T.

## Abstract

Recent studies suggest that alterations in the gut phagobiota may contribute to pathophysiological processes in mammals; however, the association of bacteriophage community structure with Parkinson’s disease (PD) has not been yet characterized. Towards this end, we used a published dataset to analyse bacteriophage composition and determine the phage/bacteria ratio in faecal samples from drug-naive PD patients and healthy participants. Our analyses revealed significant alterations in the representation of certain bacteriophages in the phagobiota of PD patients. We identified shifts of the phage/bacteria ratio in lactic acid bacteria known to produce dopamine and regulate intestinal permeability, which are major factors implicated in PD pathogenesis. Furthermore, we observed the depletion of *Lactococcus* spp. in the PD group, which was most likely due to the increase of lytic c2-like and 936-like lactococcal phages frequently present in dairy products. Our findings add bacteriophages to the list of possible factors associated with the development of PD, suggesting that gut phagobiota composition may serve as a diagnostic tool as well as a target for therapeutic intervention, which should be confirmed in further studies. Our results open a discussion on the role of environmental phages and phagobiota composition in health and disease.

## Introduction

Parkinson’s disease (PD) is the second most common neurodegenerative disease characterized by motor disturbances such as resting tremor, rigidity, postural instability, gait problems, and gastrointestinal dysfunction^1,2^. These motor symptoms are mainly related to the depletion of dopamine in the striatum as a result of a complicated multifactorial process^3^. One of the pathways leads to a loss of dopaminergic neurons in the substantia nigra pars compacta due to accumulation of fibrils of insoluble misfolded α-synuclein^4–6^.

Normally, α-synuclein plays a role in the regulation of vesicular release and is highly expressed in presynaptic neuronal terminals. The reasons why this protein adopts a β-sheet structure and forms aggregates are not completely understood; yet, the insoluble synuclein fibrils referred to as Lewy bodies are a hallmark of PD and are toxic for neurons^7^. In the Western world, the incidence of the disease is on the rise, with a higher prevalence in White men^8^. While genetic risk factors of PD, such as *SNCA* and *INPP5F* genes encoding α-synuclein and inositol polyphosphate-5-phosphatase, respectively, have been identified, it is shown that most PD cases can be attributed to environmental and epigenetic factors^9–11^. Given the overarching influence of gut bacteria on human health and early involvement of gastrointestinal (GI) microbiota in PD, the concept of the role of the microbiota-gut-brain axis in PD development has recently emerged^12,13^.

The gut bacteria may be implicated in PD through several pathways, including the effects on the enteric nervous system (ENS) which is in constant direct communication with the brain through the vagus nerve^14,15^. Furthermore, it has been suggested that changes in the gut microbial structure associated with intestinal inflammation may contribute to the initiation of α-synuclein misfolding^16,17^. The model of gut-originating, inflammation-driven PD pathogenesis is based on the idea that alterations in the intestinal bacterial community may play a role in triggering α-synuclein misfolding in the ENS. According to this model, PD starts in the ENS and then spreads in a retrograde manner through the vagus nerve to the central nervous system^16,18,19^. This concept is confirmed by the presence of α-synuclein aggregates in myenteric neurons of the ENS prior to the onset of PD motor symptoms^20^. Moreover, changes in the gut microbiota composition may cause alterations in the intestinal barrier function and permeability, affecting both the immune system and ENS, including neurons and glial cells, and exerting a profound effect on the condition of PD patients ^21–23^. Increased intestinal permeability is also associated with activation of enteric neurons and enteric glial may contribute to the initiation of alpha-synuclein misfolding^17^.

However, the factors promoting alterations of gut bacteria in neurodegenerative diseases remain unexplored. Therefore, understanding the mechanisms underlying shifts in the intestinal bacterial community that may trigger pathogenic pathways leading to PD is essential for the development of new approaches to prevent and treat this incurable disease.

The microbial community of the human GI tract is composed of bacteria, archaea, fungi, and viruses, including bacteriophages; this highly diverse and complex ecosystem is characterized by dynamic stability^24^. Bacteriophages are the most abundant group outnumbering other viral as well as bacterial species, and are considered important regulators of microbiota stability^25,26^. However, bacteriophages as possible agents that may negatively affect mammalian health have attracted scientific attention only recently^27^ We have previously shown that bacteriophage administration can cause shifts in mammalian microbiota, leading to increased intestinal permeability and triggering chronic inflammation, and have first emphasized the necessity of characterizing phagobiota, i.e., the totality of bacteriophages in humans, based on their unique characteristics distinguishing them from other viruses^28,29^.

Although it is obvious that phagobiota is an important regulator of microbial community composition in the GI and, as such, can influence the gut-brain axis, there are no data on the role of bacteriophages in neurodegenerative diseases, and the causal relationship between the microbiota changes and PD pathogenesis has never been addressed. In the past, the study of bacteriophages in humans has been limited by the lack of systematic approaches and insufficient research on phage diversity; however, the development of next-generation sequencing technologies has made metagenomic analyses of phagobiota feasible^30^.

Here, we present, for the first time, detailed comparative metagenomic analysis of intestinal phagobiota in PD patients and non-parkinsonian individuals. We retrieved a dataset of short sequence reads generated in the original study of Bedarf et al. () from NCBI Sequence Read Archive (SRA) (https://www.ncbi.nlm.nih.gov/bioproject/382085) and analysed bacterial. and bacteriophage diversity using MetaPhlAn and custom method ^23,30,31^. The obtained results revealed changes in the profile of certain bacteriophages of PD patients, thus suggesting their possible involvement in PD.

## Results

### Bacterial community composition in PD

To explore the bacterial community structure in PD, we used the shotgun metagenomics sequencing data of the faecal microbiome from 31 patients with PD and 28 control individuals^32^. HiSeq-mediated sequencing produced a total of 1,792,621,232 reads with an average of 30,383,411 reads per sample. We used the established MetaPhlAn2 method to identify abundances of bacterial taxa in the samples. We identified phage taxa and abundance by DNA sequence alignment to a database of all known phage genomes (see Methods for a detailed description). In accordance with our prior work, a threshold for detection of any phage species was used as 2 reads aligned with >90% sequence identity over more than 100 bases^33^.

In agreement with previous findings form Bedarf, J. et al, we observed differences in richness and diversity between PD and control groups (Supplementary Table S1). The richness of bacterial species in the PD microbiome tended to decrease as evidenced by lower but not statistically significant values of abundance-based coverage estimator (ACE; *p* = 0.483) and Chaol (*p* = 0.709) compared to control (Fig. 1A,B). Differences in α-diversity indexes between the PD and control groups were also not statistically significant (Shannon: *p* = 0.241; Simpson: *p* = 0. 421; inverse Simpson: *p* = 0. 428; Fig. 1C-E).

To evaluate possible differences in detail, we assessed β-diversity using the Bray-Curtis dissimilarity index and subjected the results to Principal Coordinate Analysis (PCoA), which revealed high similarity between PD and control samples based on the absence of statistically significant difference in bacterial diversity (Fig. 1F)^34,35^.

**Figure 1.**
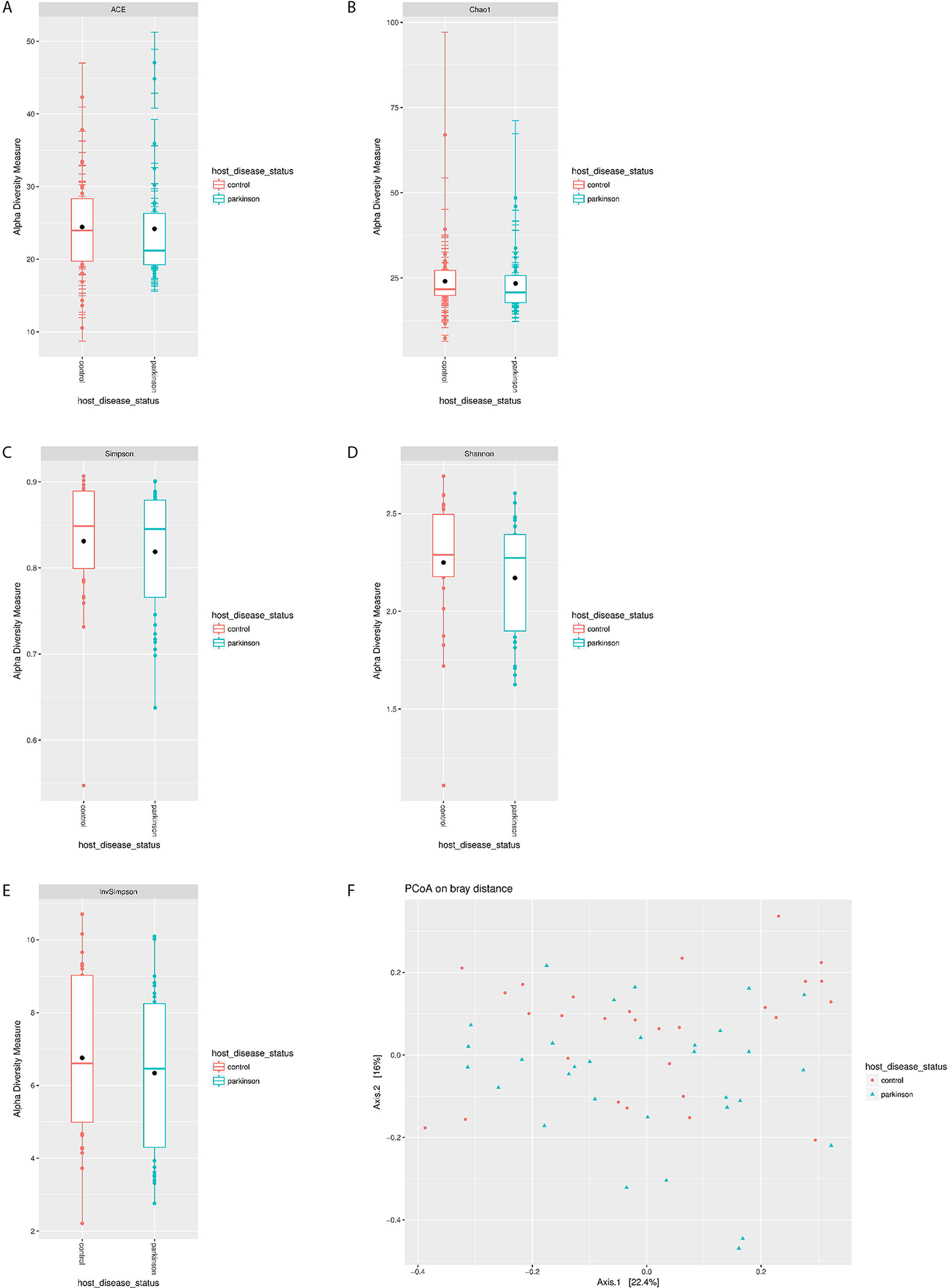
Decreased bacterial richness in the microbiome of PD patients. (A) ACE, (B) Chao1 and α-diversity, (C) Shannon, (D) Simpson, and (E) inverse Simpson indexes; *p < 0.05 compared to control, (F) PCoA plots of β-diversity in PD and control samples based on Bray-Curtis dissimilarity analyses of relative OTU composition in the samples. Each dot represents a scaled measure of the composition of a given sample, colour-and shape-coded according to the group.

The taxonomic composition of microbiome in the PD and control groups was analysed at the genus level. *Bacteroides* was the most abundant genus in both groups, and no statistically significant difference was detected in the presence of *Bifidobacterium, Eggerthella*, and *Adlercreutzia* species. However, we detected a depletion of *Prevotellaceae* and *Lachnospiraceae* species in PD patients, which is consistent with previous studies^23^. Furthermore, our analysis of less abundant families revealed reduced representation of *Lactobacillaceae* and *Streptococcaceae* in the PD group (Supplementary Table S1, Fig. 2A, B).

**Figure 2.**
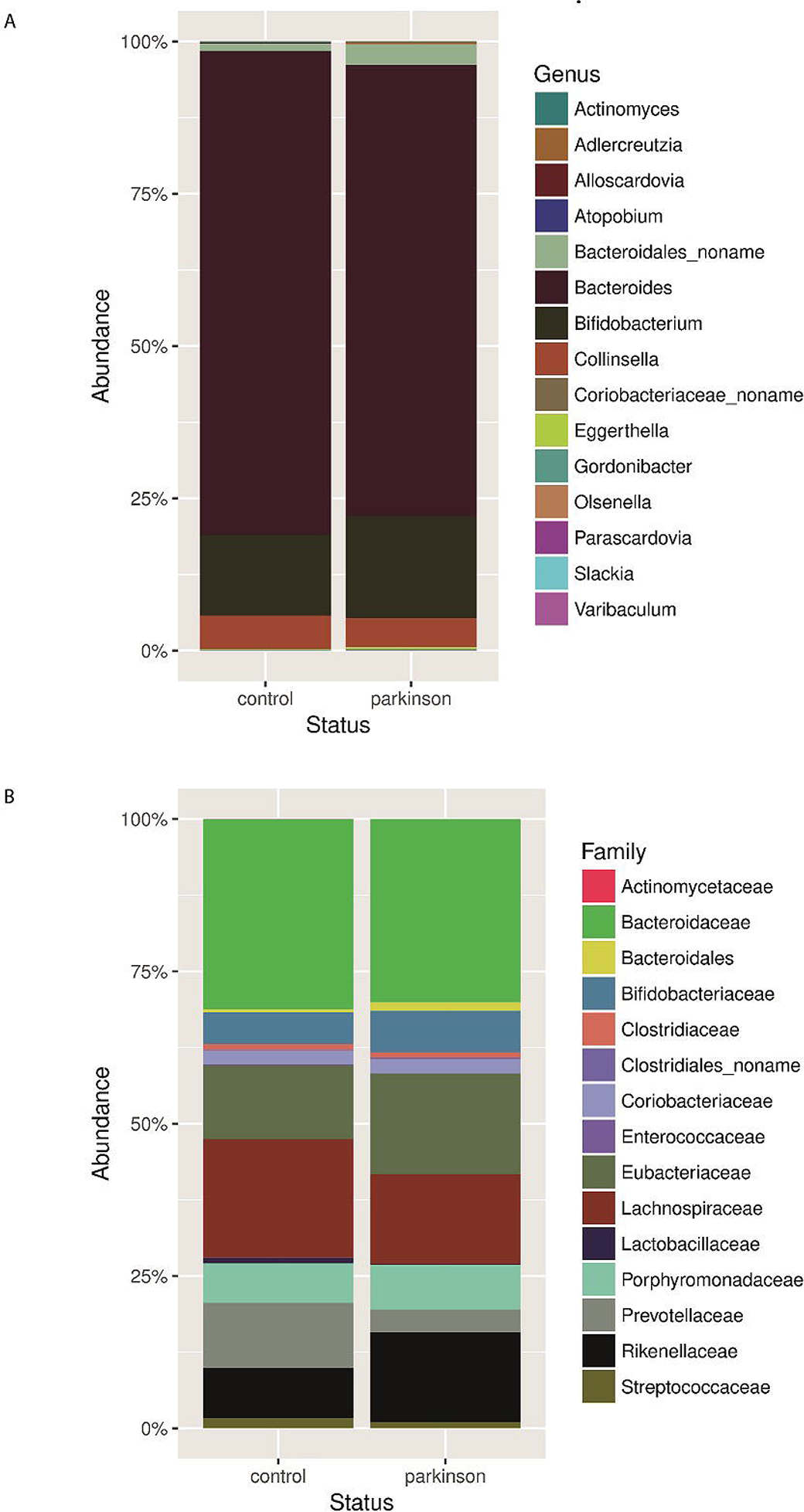
Comparison of relative abundance of predominant bacteria in PD patients and healthy participants. Faecal bacterial communities were analysed by high-throughput Illumina Hiseq4000 sequencing. Relative abundances of bacterial genera (A) and families (B) across control and PD groups.

### Bacteriophage diversity in PD

Sequence reads were then used to investigate whether there was an overall gain or loss of diversity in bacteriophage composition between the groups (Supplementary Table S2.1, S2.2, S2.3)^36^. Phagobiota richness was not statistically different between PD patients and control individuals, although a slight increase of ACE and decrease of Chao1 indexes in the PD group was detected (ACE: *p* = 0.272; Chao1: *p* = 0.797; Fig. 3A, B).

**Figure 3.**
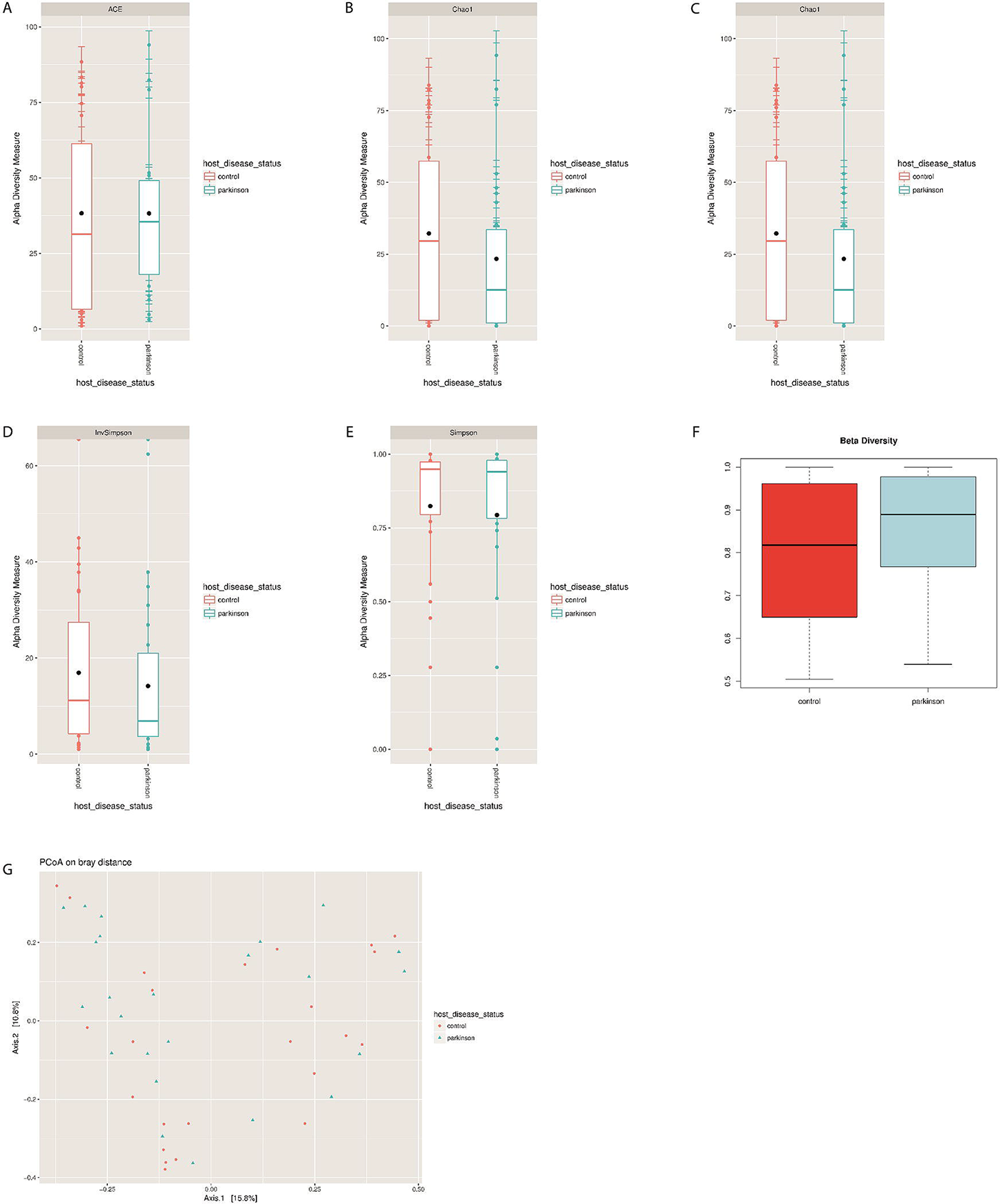
Phagobiome richness in PD patients and healthy individuals. (A) ACE, (B) Chao1 and α-diversity, (C) Shannon, (D) Simpson, and (E) inverse Simpson indexes. Bacteriophage population diversity in PD patients and healthy individuals. (F) β-Diversity of phagobiota was measured using Spearman index. The X axis indicates samples and the Y axis shows Spearman index values: 0.5 means low difference and 1 means high difference (i.e., all species are different) in species diversity between samples. (G) PCoA plots of bacteriophage β-diversity based on Bray-Curtis dissimilarity analyses. Each dot represents a scaled measure of the composition of a given sample, colour-and shape-coded according to the group.

We also found a tendency for reduction of α-diversity in PD patients indicated by reduced Shannon (*p* = 0.132), Simpson (*p* = 0.963), and inverse Simpson (*p* = 0.421) indexes (MannWhitney test) compared with control (Fig. 3C–E). These results are consistent with a similar decreasing trend observed in bacterial richness and diversity among PD patients, suggesting that the reduced numbers of bacterial hosts may be related to the reduction of the associated bacteriophages.

β-Diversity of bacteriophages was analysed based on the Spearman’s rank correlation coefficient and Bray-Curtis dissimilarity index, which revealed statistically insignificant increase in β-diversity of phagobiota in PD (Spearman’s test; p = 0.731) (Fig. 3F). Not surprisingly, PCoA of phagobiota revealed high similarity in bacteriophage diversity between PD and control groups, which showed no statistically significant difference (Fig. 3G).

### Bacteriophage composition at the family level

Analysis of relative abundance at the family level showed that *Siphoviridae* was the most abundant family in both PC and control groups (Fig. 4). The second largest bacteriophage family in the PD group was *Podoviridae* followed by *Myoviridae*, whereas in control group the order was the opposite (p = 0.017).

**Figure 4.**
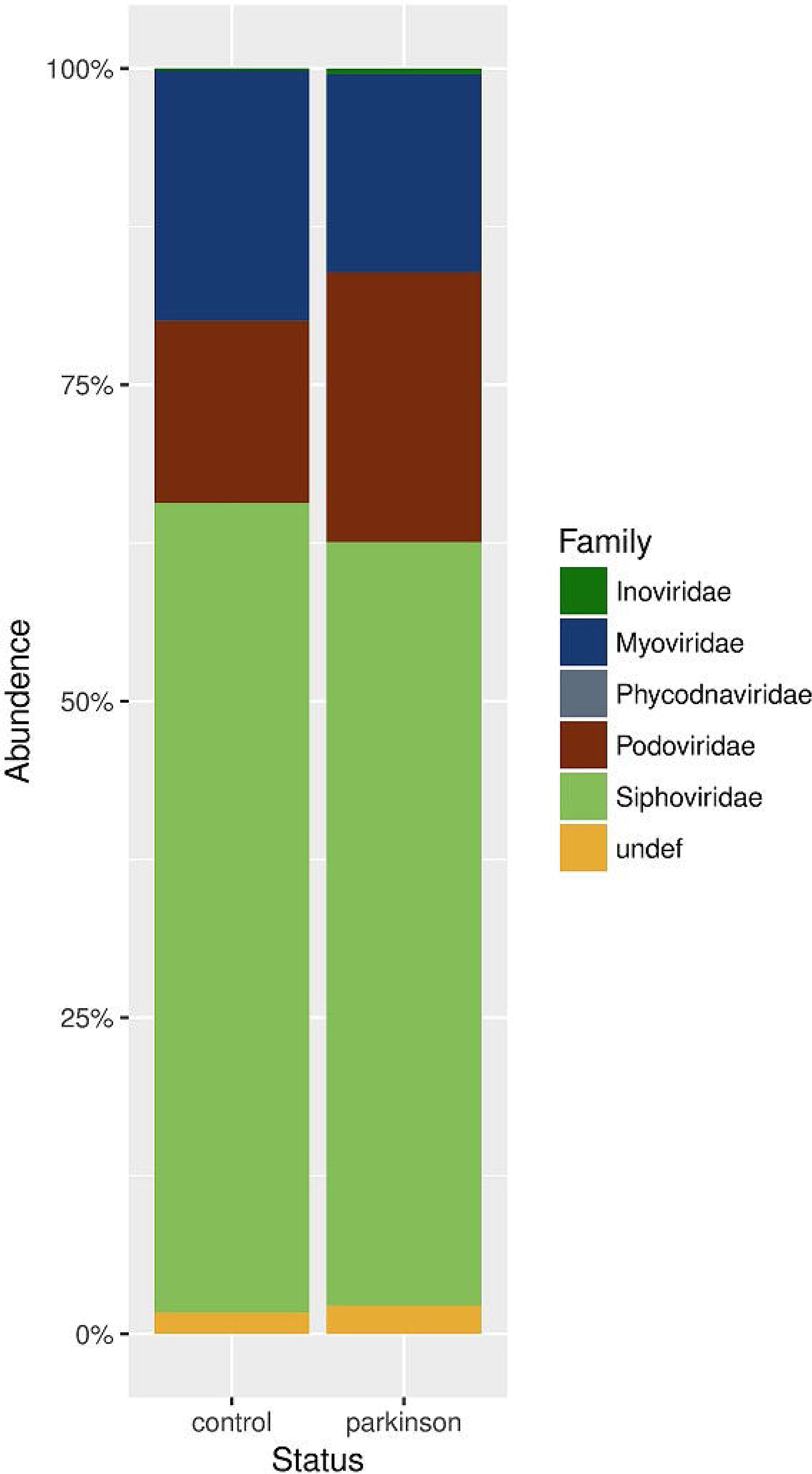
Comparison of relative abundance of predominant bacteriophage families. Faecal bacteriophage communities were analysed by high-throughput Illumina Hiseq4000 sequencing. Relative abundances of bacteriophage families in (A) control and (B) PD groups are shown.

### Composition of phagobiota at the species level

We next examined compositional changes in the bacteriophage community at the species level. To investigate phage diversity in the human gut, we analysed the abundance of phage species (relative abundance ≥ 0.01% detectable in at least two samples) individually for each sample and presented the data as a heat map (Supplementary Fig. S1). Notably, the sampling methods used in extracting total metagenomic DNA data include phages both at the lytic and lysogenic states, which cover the bulk of the phagobiota^37^. Comparative analysis revealed under-representation of phages specific for *Enterobacteria* and *Salmonella* and over-representation of those specific for *Lactococcus* (phi7, CB13, jj50, bil67, and 645) in PD patients.

Next, we evaluated species detectable only in one group (in at least two samples), which indicates appearance/disappearance of individual bacteriophage species (Fig. 5, Supplementary Table S3). The key changes included total disappearance of certain *Bacillus, Enterobacteria, Lactococcus Streptococcus*, and *Salmonella* phages, which belonged to the *Siphoviridae* family or were unclassified *Caudovirales*, as well as *Lactobacillus* phages of the *Myoviridae* family. At the same time, we detected the appearance of certain *Leuconostoc, Lactococcus*, and *Enterobacteria* phages of the *Siphoviridae* family, *Enterobacteria* phages of the *Myoviridae* family, and *Salmonella* phages of the *Podoviridae* family.

**Figure 5.**
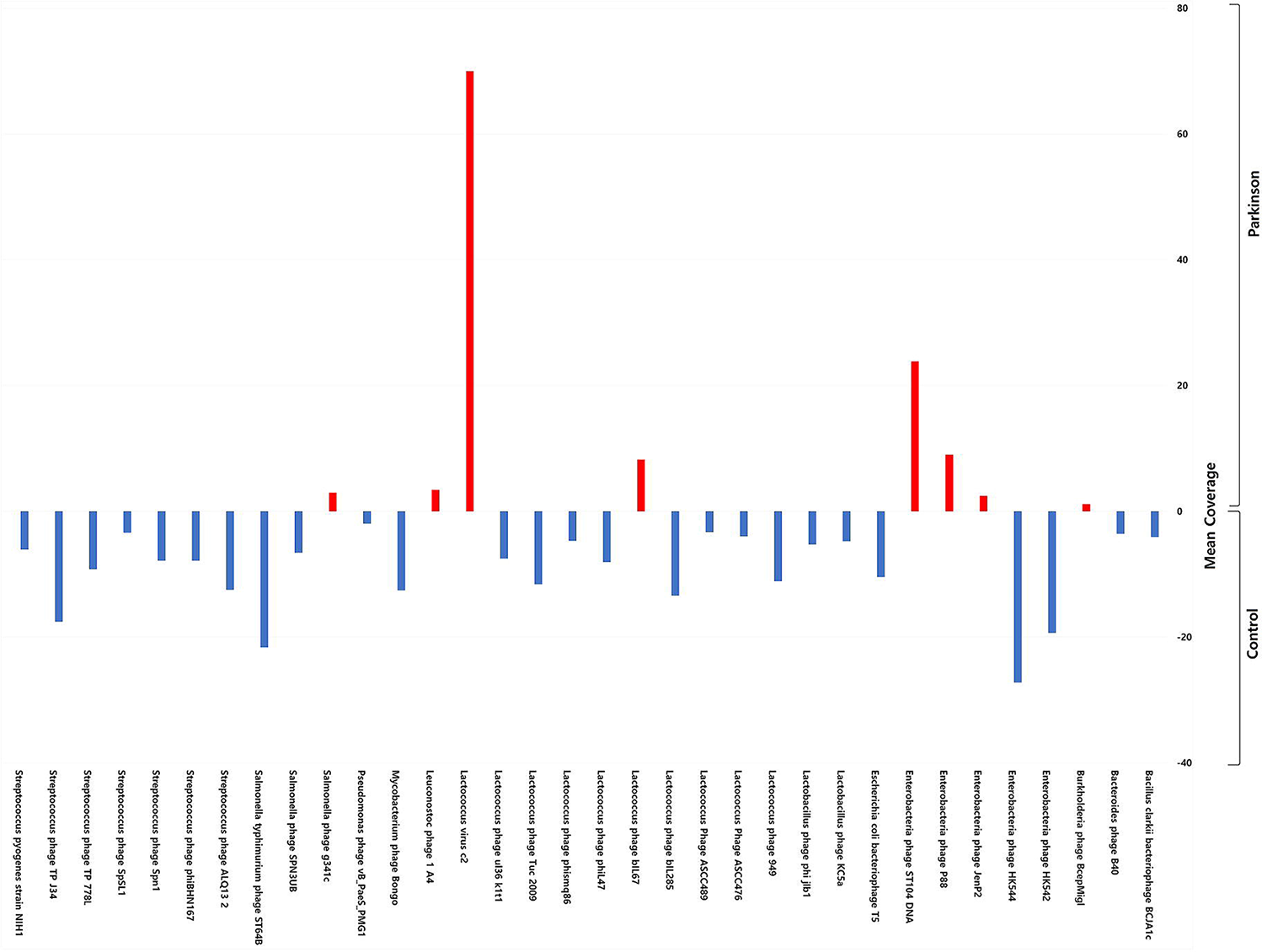
Exploration of bacteriophage diversity in PD patients and healthy participants. The bar graph shows bacteriophage abundance at the genus level in the PD or control group (relative abundance ≥ 0.01% found in at least two samples per group).

Finally, we analysed ratio between phages and their bacterial hosts in the gut microbiome by calculating the phage/bacteria ratio defined as the ‘lytic potential’^37^. To do this, we clustered phages according to their bacterial hosts and normalized phage abundance to that of the respective hosts in each group (Fig. 6). The obtained data were consistent with the trend of reduction in α- and β-diversity in both bacteriophages and bacteria among PD patients and indicated stability of the phage/bacteria equilibrium across PD and control samples. However, the identified some significant alterations in the phage/bacteria ratio in the PD group, suggesting their association with PD.

**Figure 6.**
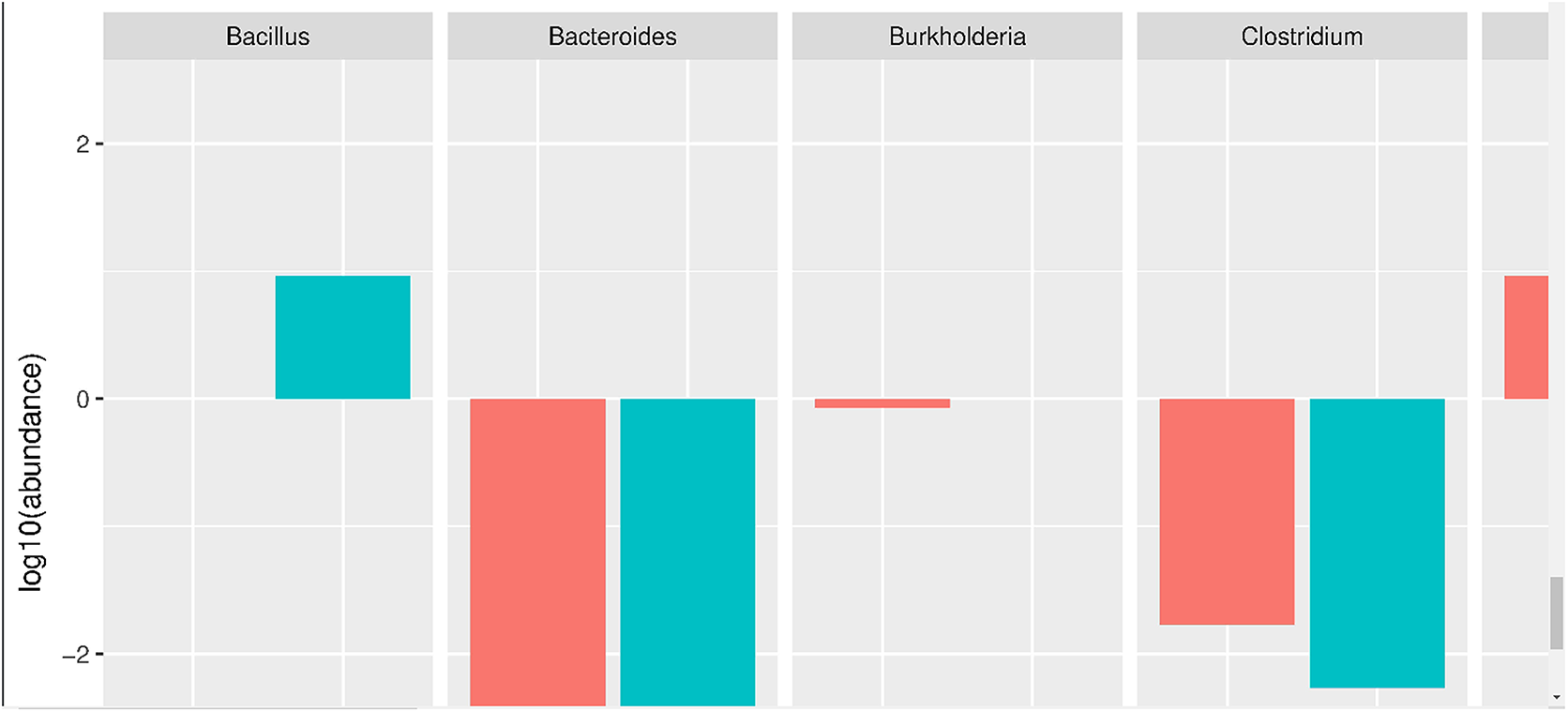
The phage/bacteria ratio in PD patients and healthy individuals. The ratio was calculated as phage abundance normalized to that of the respective bacterial hosts in each sample of the PD and control groups.

In theory, if the phage/bacteria ratio is equal to 1 (or log10°), it means that a prophage is stably integrated within the host bacterial genome, which was observed in this study for *Bacillus* and *Vibrio* phages and the corresponding bacterial host in the PD group^37^. At the same time, the ratio less than 1 indicates that a prophage is most likely absent in the genomes of a part of the corresponding bacterial host population. We observed low phage/bacteria ratios for *Bacteroides, Klebsiella, Clostridium*, and *Streptococcus*, which reflect low numbers of the corresponding phages and high abundance of the host bacteria across both PD and control groups. The phage/bacteria ratio more than 1 suggesting that the phage is at least partially present in the lytic phase, was observed for *Edwarsiella, Mycobacterium*, and *Shigella* in both PD and control groups without statistically significant difference.

However, we found a significant difference between the groups in the phage/bacterial ratio for *Lactococcus*. The abundance of *Lactococcus* spp. showed a decrease of more than 10-fold in PD patients compared to that in the control. This finding drew our attention to *Lactococcus* known to play an important role in the metabolism of neurotransmitters, including dopamine, whose deficiency is a key pathological factor in the development of PD (Fig.6)^38–41^.

We noted that despite the depletion of *Lactococcus*spp., the total number of respective *Lactococcus* phages was approximately the same between the PD and control groups (Supplementary Fig 1). To investigate this discrepancy and a possible role of bacteriophages in the depletion of *Lactococcus* spp., we divided all *Lactococcus* phages within each metagenome sample into two clusters: strictly virulent (lytic) or temperate, and compared their distribution between PD patients and controls (Fig. 7A, B)^42,43^. The results indicated that in the control group, the abundance of the lytic and temperate phages was similar, whereas in the PD group, the majority of Lactococcal phages were strictly virulent and belonged to c2-like and group 936 (Supplementary Table S4)^42^. Notably, the abundance of a majority of lytic Lacotococcus phages was higher in PD patients then in control individuals (Fig. 7B).

**Figure 7.**
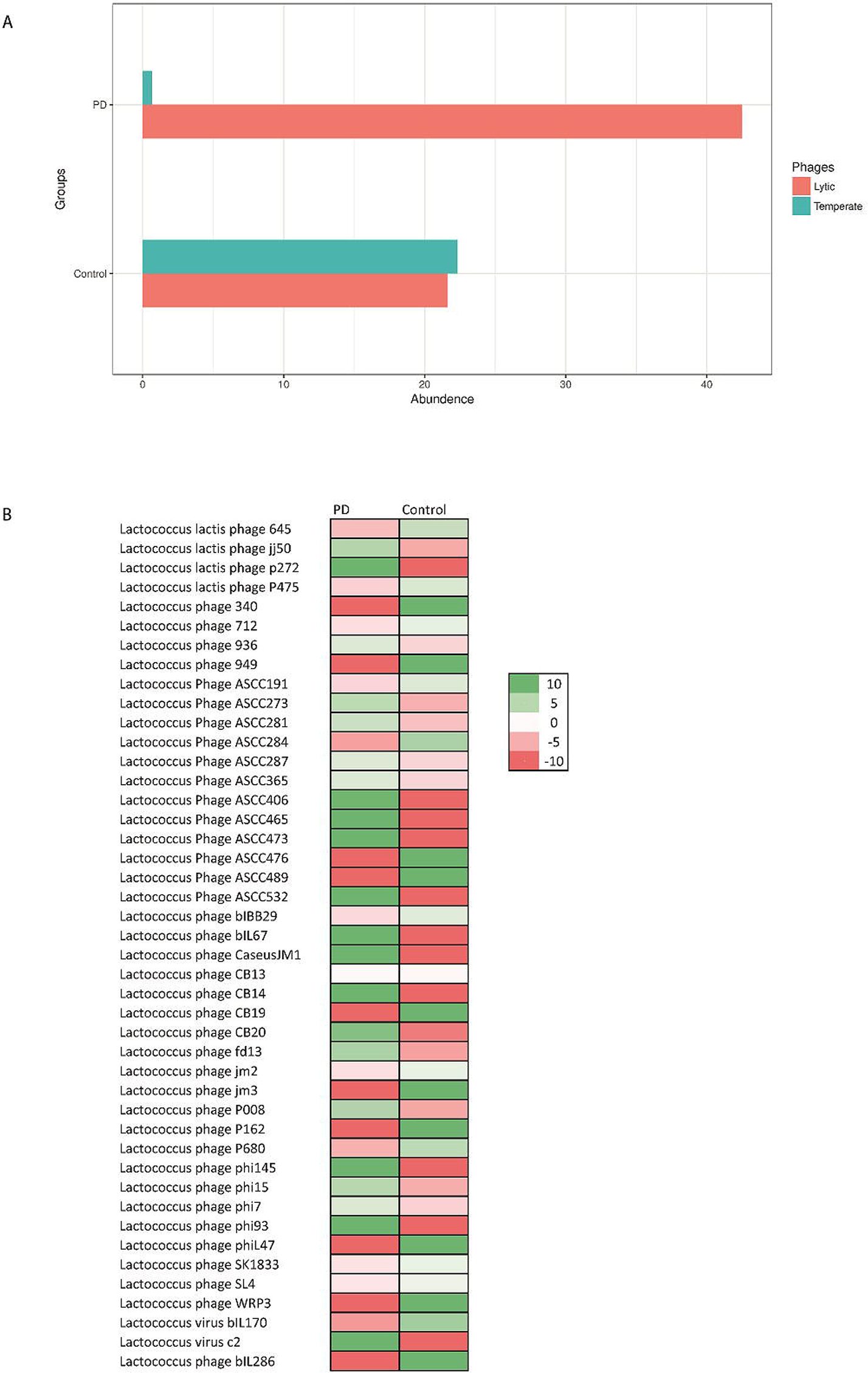
Distribution of lytic and temperate *Lactococcus* bacteriophages in the PD and control groups. (A) The graph shows the abundance of lytic and temperate lactococcus phages in each group., (B) The abundance of different lytic *Lacotococcus* phages in PD and control groups The scale on log for each row, indicates the negative values are down (Red) and positive values are up (Green) in comparison between PD and Control for each phage type.

These findings suggest that the striking depletion of *Lactococcus* spp. in PD patients could be caused by the appearance of lytic phages.

## Discussion

In this study, we have shown that the abundance of lytic *Lactococcus* phages in PD patients significantly differed from that of healthy individuals, most likely being a cause for the detected shift in neurotransmitter-producing *Lactococcus* and thus opening a discussion on the possible role of phages and implications of the phagobiota in PD.

Metagenome sequencing has greatly facilitated the investigation of the human microbiome; however, current understanding of the role of microbiota in health and disease mainly comes from the analysis of diversity and abundance of bacterial species, whereas little is known about those of bacteriophages^44^. Therefore, changes in bacteriophage composition are rarely associated with human diseases. Recently, we have shown, for the first time, that phages may be implicated in protein misfolding, altered intestinal permeability, and chronic inflammation in mammals, and, thus, should be focused on as a novel potentially critical element in the development of multifactorial diseases, including those where increased intestinal permeability is considered to be a triggering or aggravating factor^27–29^.

Here, we analysed the metagenomic data generated in the study of Bedarf et al. which included early-stage PD patients not treated with L-DOPA known to affect intestinal motility and possibly microbiota composition^23^. Our aim was to compare the phagobiota and its correlation with the bacterial component of GI microbiota between PD patients and non-parkinsonian individuals to reveal the alterations in bacteriophage composition potentially associated with the initiation or progression of PD. We identified bacteria and phages with differential abundance in the PD and control group, clustered phages according to their bacterial hosts, and evaluated the phage/bacterial ratio for each individual, which allowed us to determine whether shifts in bacterial composition resulted from phage infection or may reflect pathophysiological changes in PD patients affecting gut microbiota.

In agreement with recent studies, our findings indicate that PD is associated with lower bacterial diversity and richness^45^. Notably, our analysis revealed significant differences in the microbiota structure between PD patients and control individuals. We detected some previously overlooked alterations in the bacterial community at the family and genus levels using the MetaPhlAn tool, which provides accurate microbial profiling and estimates relative abundance of microbial cells by mapping reads against a set of clade-specific marker sequences^46^. The key alterations observed in PD patients included a decrease in certain members of *Streptococcaceae* and *Lactobacillaceae* families, such as *Lactococcus* and *Lactobacillus* (Supplementary Table S1), which are consistent with recent findings^47^. However, some studies showed different results, which may be explained by variations in analysis pipelines, bioinformatics tools, and study population, which can comprise patients at different disease stages receiving distinct therapeutic regimens that can potentially affect gut microbiota composition^45,48^.

Our attention was particularly drawn to a previously overlooked decrease of the relative abundance of *Lactococcus* spp. in the PD group. These bacteria are considered as an important source of microbiota-derived neurochemicals, including dopamine which they produce in appreciable physiological amounts^40,49^. The decrease in the production of intestinal dopamine may be, on the one hand, associated with early gastrointestinal symptoms of PD and, on the other, involved in triggering the neurodegenerative cascade of the disease^18,48^.

Moreover, lactic acid bacteria, especially lactococci and lactobacilli, are known as important regulators of gut permeability^50,51^. Although intestinal permeability of the study participants was not evaluated, previous research indicates that PD is commonly associated with impaired barrier function^52^. Therefore, given the role of lactobacteria in intestinal dopamine production, their depletion in PD patients may contribute to triggering or aggravating PD symptoms through effects on intestinal permeability^20^.

Our comparative evaluation of phagobiota composition in control individuals and PD patients using custom method did not reveal changes in α- and β-diversities, which is in agreement with the study by Bedarf et al., who showed that the abundance of prophages was not altered in the PD group, although they used a different bioinformatics tool for viral metagenome analysis^23,32^. However, we found that PD phagobiota was characterized with total disappearance of certain bacteriophage groups, including those specific for *Bacillus, Enterobacteria, Lactococcus, Streptococcus*, and *Salmonella*. At the same time, some *Leuconostoc, Lactococcus*, and *Enterobacteria* phages absent in the control group were found in a subset of PD patients.

It should be taken into account that whole genome sequencing produces results for both lytic and temperate phages, as methods used for metagenome DNA extraction isolate total phage DNA^37^. While analysing possible interplay between bacteria and phages, it is necessary to consider their mutual effects on the abundance to each other^53^. For example, high prevalence of certain prophage-harbouring bacteria should result in high numbers of the respective phages in metagenomics datasets, whereas increased abundance of certain lytic bacteriophages may be accompanied by a decrease of their bacterial hosts^54^.

With this in mind, we compared the shifts in phagobiota with the abundance of their bacterial hosts based on the phage/bacterial ratio, which allowed revealing the presence of lytic phages at the abundance higher than that of the respective bacterial hosts (Fig. 6). An important result of this analysis is the disbalance between the number of *Lactococcus* spp. and their bacteriophages in the PD group. We found more than 10-fold decrease in the abundance of *Lactococcus* among PD patients, whereas that of the respective phages was the same as in the control group. Given that the abundance of temperate phages would be reduced in parallel with their host bacteria, the finding that the decrease in *Lactococcus* spp. was not accompanied by that of the respective phages indicates that the observed reduction in *Lactococcus* may be due to the increase in the number of lytic phages.

To investigate which *Lactococcus* phages are solely lytic and which are temperate (i.e., can undergo both lysogenic and lytic cycles), we performed detailed literature analysis followed by clustering of *Lactococcus* phages into “temperate” or “lytic” groups (Supplementary Table S4)^42,55^. The results revealed that lytic *Lactococcus* phages belonging to the 936 and c2 groups were significantly overrepresented in PD patients (Fig. 7A, B), indicating that the significant decrease of the abundance of *Lactococcus* spp. in PD is accompanied by under-representation of temperate and over-representation of lytic *Lactococcus* phages. Notably, *Lactococcus* spp. are known to possess abortive infection (Abi) systems, also known as phage exclusion systems, that block phage multiplication, leading to premature bacterial death following phage infection^56^. Thus, the number of progeny phage particles decreases, limiting the spread of phages to other bacteria within the population and allowing the bacterial population to survive. It is suggested that phages in PD have developed mechanisms to overcome these antiphage systems; however, additional studies are required to confirm this^57^.

We focused on the interplay between *Lactococcus* spp. and their phages because *Lactococcus* bacteria are important dopamine producers in the ENS and regulators of gut permeability, suggesting that their depletion due to high numbers of respective phages in PD patients may be associated with PD development directly linked to dopamine decrease^51–58^.

Therefore, we can draw the second highly speculative conclusion that *Lactococcus* phages may trigger the onset of or promote PD as well as its gastrointestinal symptoms, associated with a lack of dopamine. However, additional experimental work is required to distinguish whether changes in the phagobiota can contribute to development of PD symptoms or are results of the ongoing disease ^59^. Noteworthy, There are two main scenarios that can lead to the accumulation of lytic *Lactococcus* phages in the gut and depletion of their bacterial hosts^60^. First, it can be a symptom of dysbiosis and second, a result of environmental introduction of lytic *Lactococcus* phages, which are widely used in the food industry and can be found in a variety of dairy products, including milk, cheese, and yogurt^24,58^. The impact of the latter factor is supported by the fact that the majority of lytic phages in PD patients belong to c2-like and 936- like lactococcal phages, which are most frequently isolated from dairy products^62–64^. However, the hypothesis of the role of the environmental phages in PD needs to be further explored.

In summary, our study outline the necessity to pay specific attention to bacteriophages as previously overlooked human pathogens. We identified shifts in the gut phagobiota in PD patients, some of which can be considered to be associated with the disease and may be used in the development of novel diagnostic and therapeutic tools, although this suggestion needs to be confirmed by further research.

## Methods

### Study population

Our analysis was based on the study of Bedarf et al. who recruited 31 PD patients and 28 gender-and age-matched non-parkinsonian individuals^23^. The patients had early-stage PD (onset of motor symptoms and diagnosis within the past year) not yet treated with L-DOPA known to affect gut microbiota composition. Patients with chronic and inflammatory gastrointestinal diseases, including chronic constipation, and atypical and/or secondary parkinsonism, as well as those using laxatives, immunosuppressants, or antibiotics in the past three months were excluded. Although three PD patients and three control subjects were included despite the intake of antibiotics (up to three days) in a period of 28–34 days prior to faeces sampling, the omission of those cases from the analyses had no impact on the results^23^. The demographic parameters of study participants are presented in (Supplementary Table S5)^23^.

### Microbiota sequencing and processing

Bacterial and phage content were quantified separately using the SRA shotgun metagenomic sequencing data. Bacterial content was quantified by taxa directly from SRA reads using Metaphalan (v. 2.0), which operates by mapping sequence reads to a database of predefined clade-specific marker genes^31,66^. Phage content was assessed using a custom method. First, reads from each SRA file were de novo assembled into contigs with metaSPAdes (v. 3.11.1)^67^. Then contigs >200 bp were aligned to the EBI collection of phage genomes (https://www.ebi.ac.uk/genomes/phage.html) by BLAST with a threshold e-value < 1e-5 and alignment length >50% of contig length. All of the original reads were then re-mapped with Bowtie2 (v. 2.3.4.1) to the contigs with good phage BLAST matches in order to increase sensitivity and more accurately count the abundance of reads from each type of phage. Phage read counts per contig were combined per phage genome (taxa) and normalized to relative abundance. A detection threshold of 2 reads per sample (>90% identity to the phage genome) was used, based on previous reports^33^.

Bacterial and bacteriophage communities at the genus, family, and species levels were characterized based on α- and β-diversities. All taxa with relative abundance measurements below 0.0001 in all samples, were removed from abundance tables prior to statistical analysis. α-Diversity indices (ACE, Chao 1 richness estimator, Shannon, Simpson, and inverse Simpson) were calculated using the phyloseq R library^68^. β-Diversity (similarity or difference in bacterial or bacteriophage composition between participants) was assessed based on Bray-Curtis dissimilarity computed using the “levelplot” package of the R software (https://www.r-project.org/) and represented by PCoA^34^. Differences in α- and β-diversities between datasets were examined by ANOVA and PERMANOVA statistical tests; p□<□0.05 was considered statistically significant.

### Statistical analysis of microbial community composition and differential abundance

The QIIME pipeline was used for quality filtering of bacterial and bacteriophage DNA sequences, chimera removal (by the USEARCH software), taxonomic assignment, and calculation of α-diversity, as previously described^67,68^. Downstream data analysis and calculation of diversity metrics were performed in R3.3.2 using ggplot2 and phyloseq libraries; DESeq2 was used to calculate logarithm of fold change^68^.

Bacterial and bacteriophage communities at the genus, family, and species levels were characterized based on α- and β-diversities. α-Diversity indices (ACE, Chao 1 richness estimator, Shannon, Simpson, and inverse Simpson) were calculated using the phyloseq R library^68^. β-Diversity (similarity or difference in bacterial or bacteriophage composition between participants) was assessed based on Bray-Curtis dissimilarity computed using the “levelplot” package of the R software (https://www.r-project.org/) and represented by PCoA^34^. Differences in α- and β-diversities between datasets were examined by ANOVA and PERMANOVA statistical tests; p□<□0.05 was considered statistically significant.

## Acknowledgements

We would like to thank Gregory Andronica for valuable input.

## Data availability

The other sequencing datasets generated and/or analysed during the current study are available from the corresponding author on reasonable request.

## Author Contributions

GT designed the analysis. SB and YZ conducted a metagenomics analysis. GT and VT supervised data analysis and wrote the manuscript.

## Additional Information

### Competing Interests statement

The authors declare no competing interests as defined by Nature Research, or other interests that might be perceived to influence the results and/or discussion reported in this paper.

### Publisher’s note

Springer Nature remains neutral with regard to jurisdictional claims in published maps and institutional affiliations.

